# Grk7 but not Grk1 undergoes cAMP-dependent phosphorylation in zebrafish cone photoreceptors in the dark and mediates recovery of the cone photoresponse in response to forskolin

**DOI:** 10.1101/2022.06.23.497385

**Authors:** Jared D. Chrispell, Yubin Xiong, Ellen R. Weiss

**Affiliations:** Department of Cell Biology and Physiology, The University of North Carolina at Chapel Hill, Chapel Hill, North Carolina, United States

## Abstract

In the vertebrate retina, phosphorylation of photoactivated visual pigments in rods and cones by G protein-coupled receptor kinases is essential for sustained visual function. *In vitro* analysis demonstrates that GRK1 and GRK7 are phosphorylated by PKA and that phosphorylation is associated with a decreased capacity to phosphorylate rhodopsin, while *in vivo* observations show that GRK phosphorylation is cAMP-dependent. In many vertebrates, including humans, GRK1 is expressed in both rods and cones while GRK7 is expressed only in cones. However, mice express only GRK1 in both rods and cones and lack GRK7. We recently reported that a mutation deleting the phosphorylation site, Ser21, in GRK1 is associated with a delay in dark adaptation in mouse rods but not in cones, suggesting that GRK1 may serve a different role depending upon the photoreceptor cell type in which it is expressed. Here, we present immunochemical and electrophysiological studies using zebrafish as a model. Since zebrafish display a retinal GRK expression profile similar to humans, these studies will allow us to further evaluate the role of cAMP-dependent GRK phosphorylation in cone photoreceptor recovery that may be relevant to human physiology. ERG analyses of wildtype and Grk-knockout zebrafish larvae treated with forskolin show that elevated intracellular cAMP significantly decreases recovery of the cone photoresponse, which is mediated by Grk7a rather than Grk1b. Using a cone-specific dominant negative PKA transgenic zebrafish, we show for the first time that PKA is required for Grk7a phosphorylation *in vivo*. Lastly, immunoblot analyses in rod *grk1a* and cone *grk1b* knockout zebrafish, as well as the ‘all cone’ *Nrl-/-* mouse, show that cone-expressed Grk1 does not undergo cAMP-dependent phosphorylation *in vivo*. These results provide a better understanding of the function of Grk phosphorylation in the context of cone adaptation and recovery.

## Introduction

In vertebrates, visual interpretation of the surrounding environment relies on two types of specialized neurons in the retina: rod and cone photoreceptors. Rods are highly sensitive, requiring capture of only a single photon to initiate hyperpolarization of the photoreceptor (1). However, rods are easily saturated and desensitized under conditions of prolonged brightness, thus providing vision mostly under low light conditions. Cones, while less sensitive than rods, recover more rapidly and are less prone to desensitization, which allows them to operate under a broader range of light intensities. These attributes allowing cones to function under a range of light intensities and provide higher acuity vision (2,3). Despite these differences between rods and cones, the mechanisms of phototransduction in these cells are analogous: the retinoid chromophore bound to rhodopsin or cone opsin undergoes isomerization upon absorption of a photon, which induces a conformational change in the opsin allowing the opsin to activate a G-protein (rod or cone transducin) which in turn activates a phosphodiesterase (PDE6) leading to a decrease in intracellular cGMP levels and closure of cGMP-gated cation channels in the cell membrane (4,5). Sustained vision requires that the activated opsin return to the dark-adapted state to be sensitive to new stimuli. The first step in this regeneration process is phosphorylation of the opsin which allows the binding of arrestin (6). This phosphorylation is accomplished by the retina-specific members of the G protein-coupled receptor kinase (GRK) family, GRK1 and GRK7 (7–11).

The retinal GRKs possess several qualities that govern their capacity to modulate the recovery of visual pigments to their ground state. In addition to an interaction of Grk1 with the neuronal calcium sensor protein recoverin (12–14), human recombinant Grk1 and Grk7 are phosphorylated by cAMP-dependent kinase (PKA) *in vitro* at Ser21 and Ser36, respectively (15). *In vivo* observations in several vertebrate models reveal that both GRK1 and GRK7 display elevated levels of phosphorylation at these sites under dark-adapted conditions when intracellular levels of cAMP are elevated in photoreceptors (16–21). *In vivo* phosphorylation of GRK1 was independent of phototransduction based on the observation that phosphorylation was low in the light and high in the dark in *gnat1-/-* mice (17). Additionally, *in vitro* analyses demonstrated that GRK1 and GRK7 phosphorylated at these residues have decreased kinase activity that impairs their ability to phosphorylate rhodopsin (15).

Expression of the retinal GRKs is heterogeneous in vertebrates but in many, such as humans, GRK1 is expressed in both rods and cones while GRK7 is expressed in cones (22,23). Some species, like pigs and dogs, only express Grk1 in rods and only Grk7 in cones, whereas rats and mice have lost the gene for Grk7 and express only Grk1 in both rods and cones. In humans, inactivating mutations in GRK1 results in Oguchi disease, a form of stationary night blindness characterized by slow dark adaptation (24,25). Patients with Enhanced S-Cone Syndrome (ESCS) possess L/M-cones that express only GRK1 and display a mild delay in recovery, but S-cones that lack both GRKs exhibit extremely delayed recovery (26). We previously reported that both Grk1b and Grk7a contribute to recovery of the cone photoresponse in zebrafish, which possess a retinal GRK expression pattern similar to humans, and confirm the participation of both kinases in the return of cone photoreceptors to the dark-adapted state (27). We recently reported that dark adaptation was significantly decreased *in vivo* in the rods of mice expressing GRK1 with the mutation Ser21Ala, which blocks phosphorylation (28). This suggests a role for cAMP-dependent phosphorylation of GRK1 in modulating the lifetime of activated rhodopsin. Dark adaptation was unaffected in the cones of the GRK1-S21A mice, which is surprising considering that mice express the same GRK1 protein in both rod and cone photoreceptors.

In this report, we evaluate the effects of elevated cAMP on phosphorylation of GRK1 and GRK7 in cones as well as recovery of the cone photoresponse. We incubated wildtype and Grk-knockout zebrafish larvae with forskolin, a potent activator of adenylyl cyclase, and found that increased intracellular cAMP is associated with a significant delay in cone recovery only when Grk7a is expressed in cones. We also used a cone-specific dominant negative PKA transgenic zebrafish to show that PKA is part of the endogenous kinase pathway responsible for Grk7a phosphorylation in response to elevated cAMP. Finally, we analyzed the rod/cone GRK1 phosphorylation profile of the vertebrate retina using both a newly created *grk1a-/-* zebrafish (which lacks the rod-expressed zebrafish Grk1a paralog) as well as the *Nrl-/-* mouse (which possesses an ‘all-cone’ retina) and found that GRK1 natively expressed in cones fails to undergo cAMP-dependent phosphorylation unlike GRK1 expressed in rods.

## Methods

### Materials

Mouse monoclonal antibodies against GRK1 for detection in zebrafish tissue (G8, #MA1-720) and for detection in mouse tissue (D11, #MA1-721) as well as secondary antibodies for immunoblot analysis (Alexa Fluor 680 goat anti-rabbit IgG, #A-21076) and immunocytochemistry (Alexa Fluor 488 goat anti-mouse IgG, #A-11001, and Alexa Fluor 594 goat anti-rabbit IgG, #A-11012) were purchased from ThermoFisher Scientific (Waltham, MA, USA). IRDye800 goat anti-mouse IgG (#926-32210) secondary antibody and Intercept (PBS) blocking buffer (#927-70001) were purchased from Licor Biosciences (Lincoln, NE, USA). A mouse monoclonal antibody against FLAG-tag (#F1804) was purchased from MilliporeSigma (St. Louis, MO, USA). A rabbit polyclonal against phosphorylated GRK7 was generated by Bethyl Laboratories Inc. (Montgomery, TX, USA) (16). Rabbit polyclonal antibodies against zebrafish Grk7 (27) and phosphorylated mouse Grk1 (17) were generated by 21st Century Biochemicals (Marlboro, MA, USA). A novel rabbit polyclonal antibody against phosphorylated zebrafish GRK1 was also generated by 21st Century Biochemicals using the peptide ISARG[pS]FDGTAN corresponding to amino acids 16-27 of zebrafish Grk1a. Mix-n-Stain CF Dye Antibody Labeling Kits (CF-680, #92282, and CF-770, #92285) were purchased from Biotium (San Francisco, CA, USA).

### Generation of *grk1a-/-* zebrafish line

CRISPR sgRNAs targeting exon 1 of zebrafish grk1a were designed using CHOPCHOP (https://chopchop.cbu.uib.no/) and generated based on the methods in Hwang et al. (29). Sense and anti-sense oligonucleotides corresponding to each target sequence were ordered from Integrated DNA Technologies (Newark, NJ, USA) and 1 nmol of each was annealed for 4 minutes at 95°C in annealing buffer with a final concentration of 6 mM Tris (pH 7.5), 30 mM NaCl and 600 μM EDTA. Once cooled, annealed oligos were phosphorylated *in vitro* with T4 polynucleotide kinase (#M0201S) from New England Biolabs (Ipswich, MA, USA) and ligated into pDR274 vector (#42250) from Addgene (Watertown, MA, USA), which was previously linearized with BsaI (#R0535, New England Biolabs). Following *in vitro* dephosphorylation using calf intestinal alkaline phosphatase (#M0290, New England Biolabs) the construct was purified with a Qiaquick PCR purification Kit (#28104) from Qiagen (Venlo, Netherlands). Following transformation into Dh5α cells, colony selection, and screening by Sanger sequencing (Genewiz, South Plainfield, NJ), plasmid clones containing the correct insert were linearized with DraI (#R0129S, New England Biolabs) and sgRNA was synthesized *in vitro* using a Maxiscript T7 Transcription Kit (#AM1312, ThermoFisher Scientific). Following cleanup of the sgRNAs using Micro Bio-Spin 6 columns (#7326221; Bio-Rad, Hercules, CA) and quantification, 80 pg of sgRNA was co-injected with 200 pg of Cas9 protein (#CP01-50; PNA Bio, Newbury Park, CA) into 1-cell zebrafish embryos. Genomic DNA was prepared from uninjected and injected larvae at 5 dpf and the induction of indels was measured by High Resolution Melt Analysis (HRMA). The sgRNA with the highest efficiency was 5’-GGACGTAGAGGAATACGACACGG-3’. Fish injected with this sgRNA were raised to sexual maturity (F0) and outcrossed to wildtype fish to create the F1 generation. F1 heterozygote larvae at 5 dpf were analyzed by HRMA to find an F0 parent with germline transmission and by Sanger sequencing to determine the sequence of the indel mutations. An F0 founder was identified that contained a 17 bp deletion in exon 1 of grk1a giving rise to an early termination codon at Tyr96 (pTyr96*). Additional F1 progeny from this fish were raised to sexual maturity and fin-clipped to find individuals carrying the pTyr96* allele. Once identified, heterozygous pTyr96* F1 fish were in-crossed to produce F2 offspring. These F2 fish were raised to adulthood and fin-clipped for HRMA and Sanger sequencing to identify homozygous pTyr96* fish which were used to continue the line, hereafter designated *grk1a-/-*.

### Generation of a PKA dominant negative *Tg(gnat2:prkar1a*^*G323D*^) zebrafish line

The dominant negative PKA RIαB mutant was originally generated by a G324D mutation in the *Prkar1a* gene encoding the RIα regulatory subunit of PKA in mice (30,31). This glycine is conserved in the zebrafish ortholog *prkar1aa*, at position 323. After cloning *prkar1aa* from a zebrafish retina cDNA library and inserting into pGemT-Easy vector (#A1360) from Promega (Madison, WI), site-directed mutagenesis was performed as described in Liu & Naismith (32) using primers 5’-ACTACTTCGATGAGATCGCTCTGCTCATGAACCGTCCTCGTGCTG and 5’-AGCGATCTCATCGAAGTAGTCAGACGGTGCGAGTCTTCCAACCTC using Phusion Plus DNA polymerase (#F630S, ThermoFisher). Following confirmation of the RIαB G323D mutation by Sanger sequencing, the zebrafish RIαB fragment was cloned into the pME-MCS (#237) middle entry vector of the tol2kit (33). Simultaneously, a 3.2-kb promoter fragment (TαCP) from cone transducin alpha (*gnat2*) (34) was cloned into p5E-MCS 5’ entry vector (#228). Gateway cloning was performed using LR Clonase II Plus enzyme mix (#12538200, ThermoFisher) with these two entry clones plus the p3E-polyA (#302) 3’ entry vector and the pDestTol2CG2 (#395) destination vector. Successful recombination of the TαCP-RIαB-polyA-pDestTol2CG2 plasmid was confirmed by Sanger sequencing of transformed clones. Capped Tol2 transposase mRNA was synthesized *in vitro* using mMessage mMachine SP6 Transcription Kit (#AM1340, ThermoFisher) with NotI-linearized pCS2FA-transposase (#396) plasmid as a template. The TαCP-RIαB-polyA-pDestTol2CG2 plasmid (30 pg) and Tol2 transposase mRNA (25 pg) were co-injected into zebrafish embryos at the 1-cell stage. Live injected embryos were screened using fluorescence microscopy at 4 dpf to identify EGFP mosaic expression in cardiac muscle, as the pDestTol2CG2 destination vector also carries *cmlc2*:EGFP as a transgenesis marker. EGFP-positive larvae were raised to sexual maturity (F0) and outcrossed to wildtype fish to produce F1 zebrafish, which were similarly screened for heart EGFP expression to identify germline transmitting F0 founders. F1 zebrafish and subsequent generations that were EGFP-positive were used in these studies. Animals from this line are designated *Tg(gnat2:prkar1a*^*G323D*^).

### Immunoblotting

Unless noted, all tissue samples for immunoblot analysis were collected in HEPES-Ringer buffer containing 10 mM HEPES, pH 7.5, 120 mM sodium chloride, 0.5 mM potassium chloride, 0.2 mM calcium chloride, 0.2 mM magnesium chloride, 0.1 mM EDTA, 10 mM glucose, and 1 mM DTT (35). Hot (95°C) 2x Laemmli buffer (125 mM Tris-HCl pH 6.8, 4% SDS [w/v], 20% glycerol [v/v]), without bromophenol blue or β-mercaptoethanol but containing 100 μM NaF to block protein dephosphorylation, was added to a final concentration of 1.2x (36). Samples were homogenized briefly with a motorized pestle, heated for 5 min at 95°C, sonicated at low power for 5 seconds to shear genomic DNA, heated again for 2 min at 95°C, and the supernatant collected following centrifugation at 10,000xg for 7 minutes. Protein concentration was determined using a DC Protein Assay (#5000112) from Bio-Rad (Hercules, CA, USA). Following addition of bromphenol blue and β-mercaptoethanol (final concentrations 0.01% [w/v] and 5% [v/v], respectively), homogenates were analyzed by either 8% or 10% SDS-PAGE, followed by immunoblot analysis. Nitrocellulose membranes were blocked in Intercept (PBS) Blocking Buffer (#927-70001, Licor Biosciences) and incubated overnight at 4°C in Intercept Buffer with 0.1% Tween 20 (v/v) and primary antibodies. Blots were washed four times for 5 minutes each in 1x PBS containing 0.1% Tween 20, then incubated in Intercept Buffer containing 0.1% Tween 20 and secondary antibodies for 1 hour at room temperature. Following 4 additional washes in 1x PBS containing 0.1% Tween 20 for 5 minutes each and 1 wash in 1x PBS for 5 minutes, blots were analyzed using the Odyssey Infrared Imaging System (Licor Biosciences). All primary antibodies were used at a dilution of 1:10,000 except for anti-phosGrk7 (1:2000), and all secondary antibodies were used at a dilution of 1:15,000. For experiments requiring the use of two antibodies derived from the same host species, primary antibodies were directly conjugated to fluorophores using Mix-n-Stain CF Dye Antibody Labeling Kits (Biotium) according to the manufacturer’s instructions.

To determine levels of phosphorylation of retinal GRKs, tissue preparations varied by species and age. For zebrafish adults, both retinas were harvested and placed in 100 μl of HEPES-Ringer buffer to which 150 μl of hot (95°C) 2x Laemmli buffer was added, then processed as described above. Following protein quantification, 25 μg of total protein per adult zebrafish retina sample was run in each lane. For 5 dpf zebrafish larvae, whole intact eyes from 20 larvae were harvested and placed in 16 μl of ice-cold HEPES-Ringer buffer to which 24 μl of hot (95°C) 2x Laemmli buffer was added, then processed as described above. An entire zebrafish larval eye sample (40 eyes) was run in each lane. To determine the immunoreactivity of the novel anti-phosphoGrk1 antibody, recombinant FLAG-tagged zebrafish Grk1a and Grk1b were each inserted into the pFLAG-CMV2 vector (#E7033, Millipore-Sigma). HEK-293 cells cultured in DME/F-12 with 10% fetal bovine serum were transfected with either of these constructs using FuGENE 6 Transfection Reagent (#E2691) from Promega (Madison, WI, USA). Approximately 72 hours after transfection, cells were harvested, frozen once, then homogenized in Tris-buffered saline containing 0.5% n-dodecylmaltoside and Complete Protease Inhibitor Cocktail (#11836170001, Millipore-Sigma), followed by centrifugation at 40,000xg for 30 minutes at 4°C. FLAG-tagged zebrafish Grk1a or Grk1b was isolated using anti-FLAG M2 Affinity Gel (#A2220) from Millipore-Sigma as described previously (15) *In vitro* phosphorylation or dephosphorylation reactions were carried out by incubating FLAG-purified Grk1a or Grk1b with PKAα catalytic subunit (#P6000S) or λ phosphatase (P0753S) from New England Biolabs (Ispwich, MA). Reactions were stopped by the addition of 2x Laemmli Buffer to a final concentration of 1.2x then subjected to immunoblot analysis as described above.

### Electroretinogram (ERG) Analysis

ERG analysis was performed as described previously (27,37,38). Zebrafish larvae were dark-adapted overnight prior to analysis. All larvae were incubated for 5 minutes under indirect dim white light (<1 lux) in zebrafish system water containing 500 μM 2-amino-4-phosphonobutyric acid (APB; DL-AP4, #0101) from Bio-Techne/Tocris (Minneapolis, MN), an mGluR6 agonist that blocks signaling to ON bipolar cells and allows measurement of the isolated cone mass receptor potential. Larvae were anesthetized with 0.02% Tricaine, positioned under low light (275 lux for ≤1 min) on a piece of filter paper and covered with 3% methylcellulose posterior to the eye. Specimens on filter paper were then transferred to a damp sponge saturated with oxygenated Goldfish Ringer’s buffer (39) on a modified stage underneath a Colordome light stimulator (Diagnosys LLC, Lowell, MA, USA). The entire ERG apparatus was located inside a black curtained, copper mesh lined Faraday cage. The microelectrode, a chlorided silver wire in a glass capillary with a 10 μm tip opening filled with Goldfish Ringer’s buffer, was positioned onto the central cornea. An AgCl pellet underneath the sponge served as a reference electrode. Once positioned, larvae were allowed to dark-adapt for 5 minutes while monitored with a combination of indirect infrared lighting, an infrared camera and a red-filtered video monitor placed outside the cage. Recordings were obtained using an Espion E2 system (Diagnosys LLC). The sensitivity of the cone mass receptor potential was determined using a single flash paradigm consisting of 20-ms flashes of white LED light with intensities of 1, 5, 10, 20, 50, 100, 500, 1000, 2000, and 5000 cd/m^2^. Recovery to successive stimuli was determined using a dual flash paradigm that utilized two 20-ms flashes of white LED light with an intensity of 1000 cd/m^2^ separated by variable interstimulus intervals (ISIs) of darkness.

### Light/Dark Adaptation

Animals indicated as ‘dark-adapted’ were dark-adapted overnight for a minimum of 16 hours. All dissections and sample preparations were performed under oblique dim red or infrared light until after the sample was homogenized and heated as described above. Animals indicated as ‘light-adapted’ were dark-adapted overnight, then placed under white light of 1200 lux for at least 2 hours. In the experiment involving the *Tg(gnat2:prkar1a*^*G323D*^) transgenic larvae, animals were dark-adapted overnight, light-adapted under white light (1200 lux) for at least 2 hours, then dark-adapted for 15 minutes prior to whole eye collection under oblique dim red or infrared light. The samples were then processed for immunoblot analysis as detailed above.

### Forskolin Treatments

For immunoblot analysis, larvae at 5 dpf were incubated in system water for 30 minutes with either forskolin (50 μM) or vehicle DMSO (0.5%, v/v). For ERG analyses, larvae were incubated in forskolin or vehicle for 25 minutes followed by a 5 min co-incubation with APB (500 μM).

### Statistical Analyses

For single flash ERGs, normalized peak amplitude responses were fit using the Naka Rushton function with GraphPad Prism (La Jolla, CA, USA) (40). For dual flash ERGs, we fit linear mixed models of covariance to compare response recovery between drug treatments across repeated ISI levels. Models included random fish effects to account for within-fish correlations. These analyses were performed using JMP (SAS, Cary, NC, USA). For immunoblots, statistical comparison of multiple groups was performed using a two-way analysis of variance (ANOVA) followed by a Tukey post hoc test. Comparisons in Fig. 5 were performed using a Student’s *t* test. For all analyses, a P-value of 0.05 was considered significant.

## Results

### Zebrafish Grk1 is phosphorylated in vitro by PKA and in dark-adapted zebrafish

Human recombinant Grk1 and Grk7 undergo phosphorylation by PKAα *in vitro* at Ser21 and Ser36, respectively (15). Mouse GRK1 phosphorylation under dark-adapted conditions in a cAMP-dependent manner *in vivo* is detected using an antibody specific for GRK1 phosphorylated at Ser21 (17). To determine if these kinases are phosphorylated in zebrafish, an antibody was generated against a peptide sequence corresponding to phospho-Ser21 in zebrafish Grk1 that is conserved in the Grk1a and Grk1b paralogs. To examine the immunospecificity of this antibody, recombinant FLAG-tagged Grk1a and Grk1b were expressed in HEK-293 cells and immunopurified with anti-FLAG beads. Purified FLAG-Grk1a and FLAG-Grk1b were then incubated *in vitro* with either λ phosphatase or PKAα catalytic subunit and subjected to immunoblot analysis using the novel anti-phosGrk1 antibody and the anti-FLAG antibody. The anti-phosGrk1 antibody recognizes both recombinant Grk1a and Grk1b phosphorylated by the PKAα catalytic subunit *in vitro* (Figure 1A). Additionally, this antibody recognizes phosphorylated Grk1 in retinal homogenates from dark-adapted adult zebrafish (Figure 1B). These results demonstrate that the anti-phosGrk1 antibody recognizes cAMP-dependent Grk1 phosphorylation in zebrafish but does not discriminate between the Grk1 paralogs.

**Figure 1.**
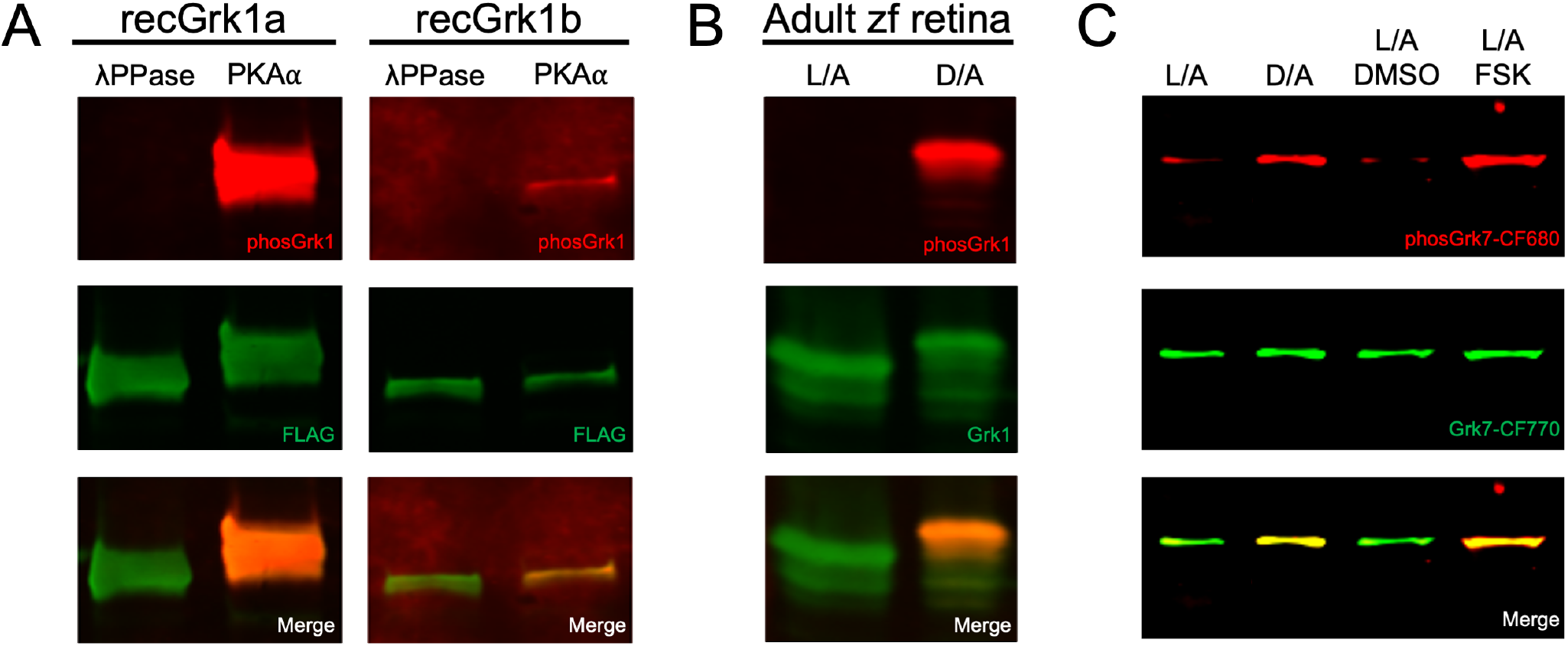
Immunospecificity of antibodies against phosphorylated Grk1 and Grk7. A) FLAG-tagged, purified recombinant zebrafish Grk1a and Grk1b treated *in vitro* with either λ phosphatase or PKAα catalytic subunit were subjected to immunoblot analysis. Immunoblots were probed with antibodies against the FLAG-tag (*green*) and phosphorylated Grk1 (*red*) followed by incubation with secondary antibodies. B) Adult zebrafish were light- or dark-adapted, euthanized, and retinal homogenates subjected to immunoblot analysis. Immunoblots were probed with antibodies against Grk1 (*green*) and phosphorylated Grk1 (*red*) followed by incubation with secondary antibodies. C) Larvae at 5 dpf were light- or dark-adapted, followed by incubation with forskolin or DMSO for 30 minutes. Larvae were euthanized and intact eyes were harvested for immunoblot analysis. Immunoblots were probed against Grk7 (*green*) and phosphorylated Grk7 (red) with anti-Grk7 antibody directly conjugated to CF770 and anti-phosphorylated Grk7 antibody directly conjugated to CF680 (*red*).

The antibody against Grk7 was reported to recognize both paralogs of Grk7 in zebrafish, while the antibody against phosphorylated Grk7 showed cAMP-dependent phosphorylation of GRK7 at Ser36 (Ser33 in zebrafish) *in vivo* in multiple vertebrates via immunohistochemical analysis (16,27). Since both antibodies are rabbit-derived, they were directly conjugated to distinct fluorophores. Immunoblot analysis confirms colocalization of bands corresponding to phosphorylated and total Grk7 in retinas of both dark-adapted and forskolin-treated larvae at 5 dpf (Figure 1C).

### Both Grk1 and Grk7 are phosphorylated in retinas of zebrafish larvae incubated with forskolin

Recombinant human GRK1 and GRK7 undergo phosphorylation in HEK-293 cells treated with forskolin, an activator of adenylyl cyclase that has been observed to increase intracellular cAMP in vertebrate photoreceptors (15,41,42). Similarly, frog retinas incubated in forskolin *ex vivo* displayed an increase in Grk7 phosphorylation (16). To determine the *in vivo* effects of increased cAMP on Grk phosphorylation in the zebrafish retina, larval zebrafish were incubated in forskolin under both light-and dark-adapted conditions prior to immunoblot analysis. Quantification of immunoblots show a significant increase in Grk1 phosphorylation under dark-adapted conditions compared to light adapted conditions regardless of drug treatment and in dark-adapted larvae treated with forskolin compared to vehicle (Figure 2a). We observed a six-fold increase in Grk1 phosphorylation in light-adapted larvae treated with forskolin compared to those treated with vehicle and a phosphorylation-dependent electrophoretic mobility shift in Grk1 immunostaining in these larvae compared to light-adapted, vehicle-treated larvae. Immunoblot analysis and quantification also show a significant increase in Grk7 phosphorylation in larvae exposed to forskolin compared to vehicle regardless of background illumination (Figure 2B). A significant increase in Grk7 phosphorylation was observed in forskolin-treated larvae under dark-adapted conditions compared to light-adapted, and Grk7 phosphorylation was elevated 10-fold in vehicle-treated larvae under dark-adapted conditions compared to light-adapted. These data suggest that under conditions where intracellular cAMP is elevated *in vivo*, both retinal Grks undergo phosphorylation.

**Figure 2.**
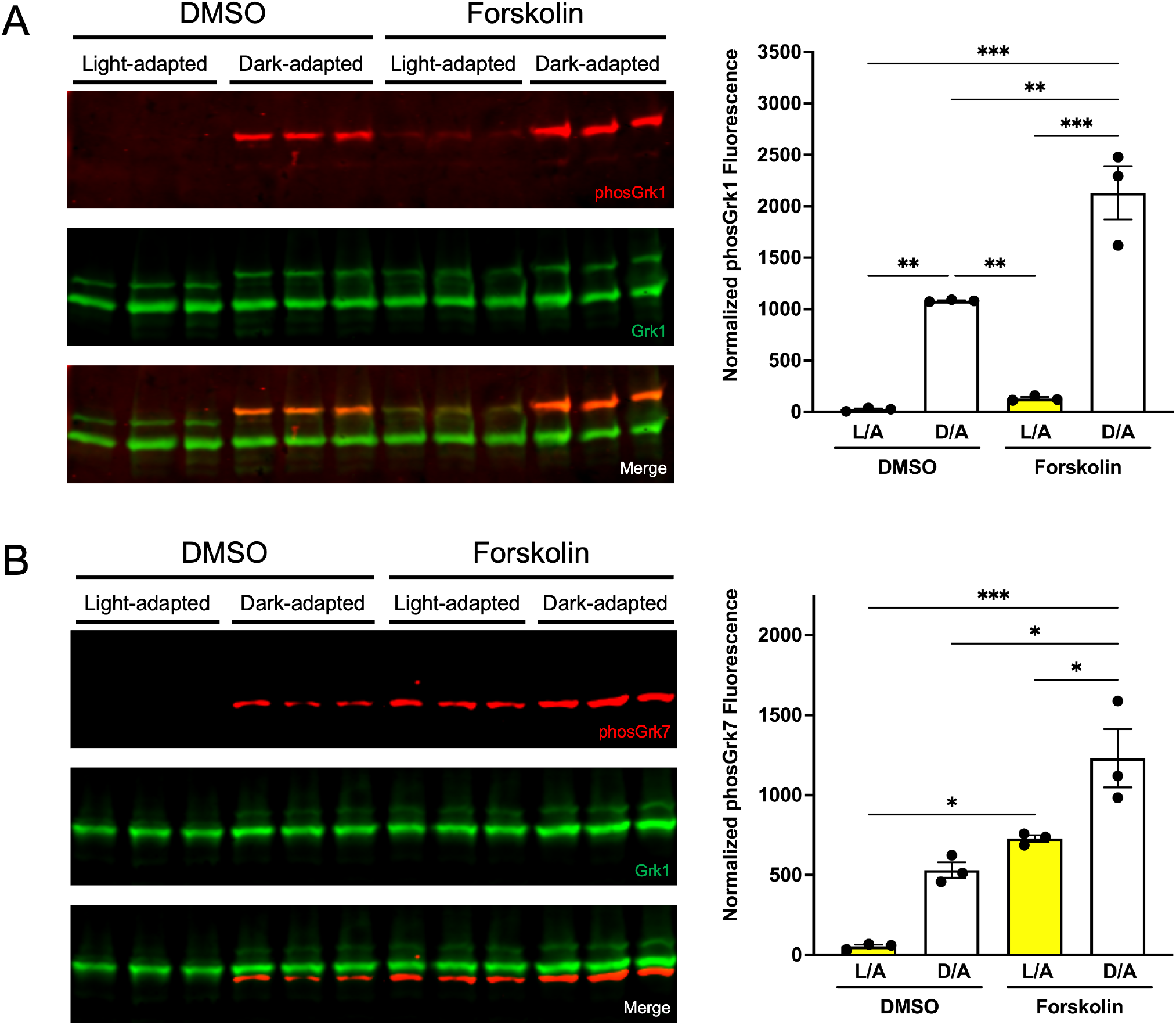
Effect of light exposure and forskolin incubation on phosphorylation of retinal GRKs in wildtype zebrafish larvae at 5 dpf. Larvae were light- or dark-adapted, followed by incubation with forskolin or DMSO for 30 minutes. Larvae were euthanized and intact eyes were harvested for immunoblot analysis. Immunoblots were probed with antibodies against total Grk1 (*green*) and either A) phosphorylated Grk1 (*red*) or B) phosphorylated Grk7 (*red*) followed by incubation with secondary antibodies. The levels of phosphorylated Grk1 or Grk7 were normalized against total Grk1 and quantified. Statistical comparison of multiple groups was performed using a two-way analysis of variance (ANOVA) followed by a Tukey post hoc test. Error bars represent SEM (n=3). *P ≤ 0*.*05 (*), P ≤ 0*.*01(**), P ≤ 0*.*001(***)*.

### Recovery of the cone photoresponse is significantly delayed in zebrafish larvae exposed to forskolin

To understand the effects of increased intracellular cAMP on the cone photoresponse, electroretinogram (ERG) recordings were performed on larvae incubated with forskolin. In larval zebrafish at 5 dpf, cones are electrophysiologically functional while nascent rods are not. The native photopic response at 5 dpf consists of a small a-wave elicited by the cone photoreceptors that is occluded by the larger b-wave response of the inner retina. Isolation of the cone mass receptor potential for recording was achieved through co-incubation of larvae with APB, an mGluR6 agonist, that blocks neurotransmission to ON bipolar cells and eliminates the b-wave. Responses were recorded in sedated larvae across an intact eye to single 20-ms flashes of white light of intensities from 0.1 to 5000 cd/m^2^ (Fig. 3A, left panels). To compare cone sensitivity in wildtype larvae treated with either vehicle (DMSO) or forskolin, responses were normalized (μV/μV_Max_) and fit with the Naka-Rushton equation (Fig. 3A, right panel). No significant differences in peak amplitudes or normalized sensitivity were observed between vehicle- or forskolin-treated wildtype larvae.

**Figure 3.**
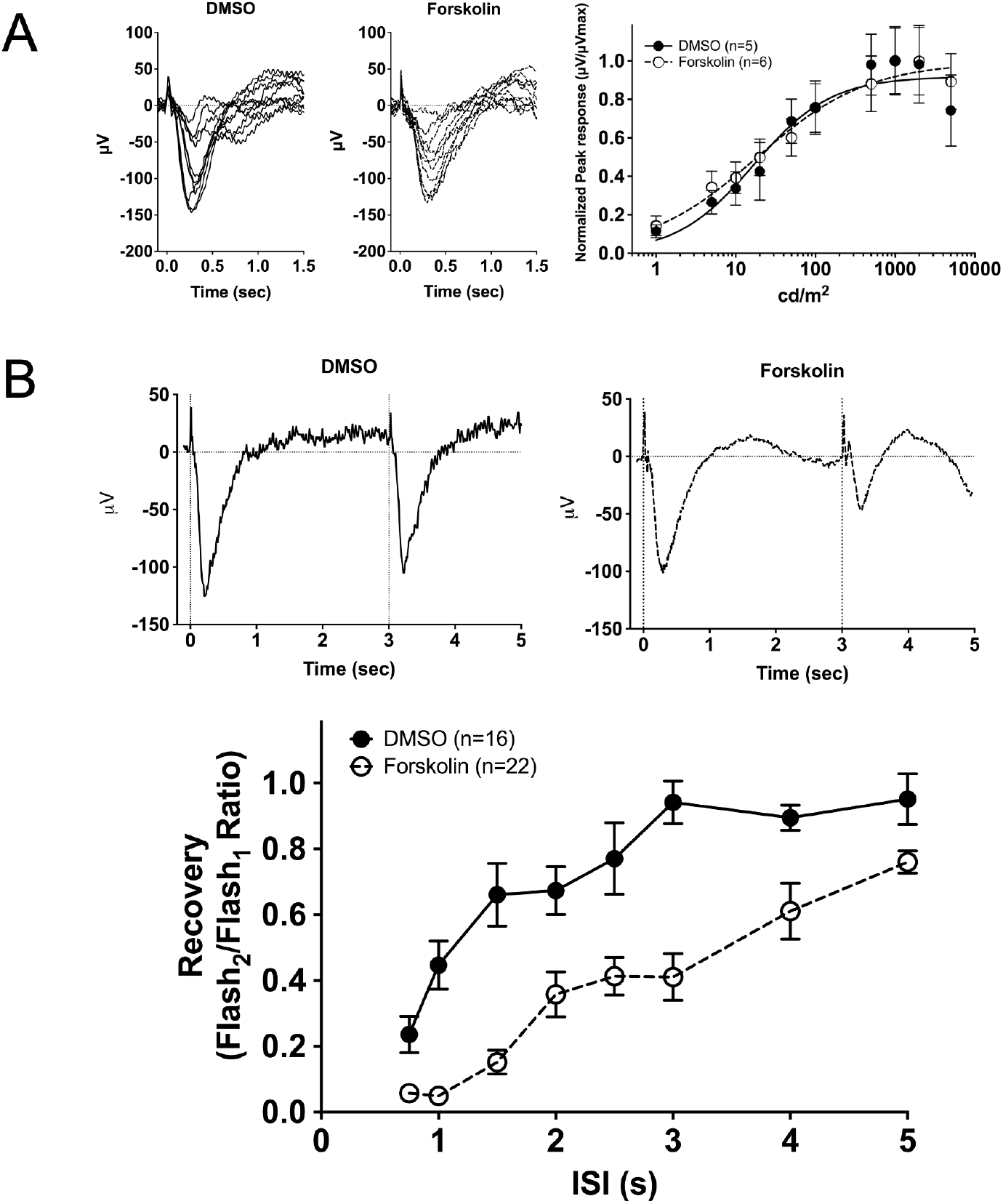
Electrophysiological light responses, normalized sensitivity, and recovery of the cone-mass receptor potential in forskolin-treated wildtype larvae at 5 dpf. A) Representative ERG traces of isolated cone-mass receptor potentials in larvae incubated in vehicle (DMSO) or forskolin for 25 min, followed by 5 min of co-incubation with APB. Reponses were recorded under dark-adapted conditions to 20-ms flashes of light of increasing intensities equal to from 0.1 cd/m^2^ to 5000 cd/m^2^. The fast initial positive deflection is attributed to a photovoltaic effect with the recording microelectrode. Mean-normalized peak response amplitudes were fit using the Naka-Rushton function. B) Representative ERG waveforms of treated larvae subjected to successive stimuli using a dual flash paradigm of a 20-ms flash of saturating white light (1000 cd/m^2^) with an interstimulus interval (ISI) of 3 s. Vertical black dotted lines indicate time of stimulus. Cone-mass receptor potential recovery was plotted as the ratio of the maximum isolated cone mass receptor potential response of the second stimulus to that of the initial stimulus for ISIs ranging from 0.75 s to 5 s. A linear mixed model analysis of covariance found a significant effect of forskolin when compared to DMSO [F_(1, 350)_ = 115.6; p<0.0001]. Error bars represent SEM.

To determine if exposure to forskolin affects the cone photoresponse recovery, ERG responses to successive stimuli with a variable interstimulus intervals (ISI) were analyzed. Vehicle- or forskolin-treated larvae wildtype larvae were subjected to a conditioning flash of saturating white light (1000 cd/m^2^, 20 ms), followed by a probe flash of the same intensity at varying ISIs. Recovery was measured as the ratio of the cone mass receptor potential of the probe flash to the initial conditioning flash. Representative recovery waveforms using an ISI of 3 seconds are shown in Figure 3B (top panels). A time course of the cone mass receptor potential recovery shows a statistically significant delay in recovery for forskolin-treated larvae compared to vehicle (DMSO)-treated larvae [F_(1, 350)_ = 115.6; p<0.0001] (Figure 3B, bottom panel). This indicates that increasing levels of intracellular cAMP via incubation of larvae with forskolin leads to impairment of the cone photoresponse.

### Zebrafish larvae that lack Grk7a are insensitive to the forskolin-induced delay in recovery of the cone photoresponse

We previously reported that both Grk1b and Grk7a contribute to the recovery of the cone photoresponse based on observations made using zebrafish lines in which either of the cone-specific Grk paralogs, Grk1b or Grk7a, was deleted through the use of TALENs (27). To determine if one or both GRK paralogs mediate the delayed cone recovery observed when intracellular cAMP levels are elevated, paired flash ERG analysis was performed on *grk1b-/-* and *grk7a-/-* larvae treated with forskolin. In *grk1b-/-* larvae treated with forskolin, we observed a significant delay in recovery compared to *grk1b-/-* treated with vehicle [F_(1, 224)_ = 17.82; p <0.0001] (Figure 4A). However, *grk7a-/-* larvae treated with forskolin displayed no significant change in cone recovery compared to vehicle-treated *grk7a-/-* larvae [F_(1, 340)_ = 2.363; p=0.1252] (Figure 4B). These data suggest that the delay in cone recovery brought about by increased levels of cAMP is mediated by Grk7a in zebrafish larvae.

**Figure 4.**
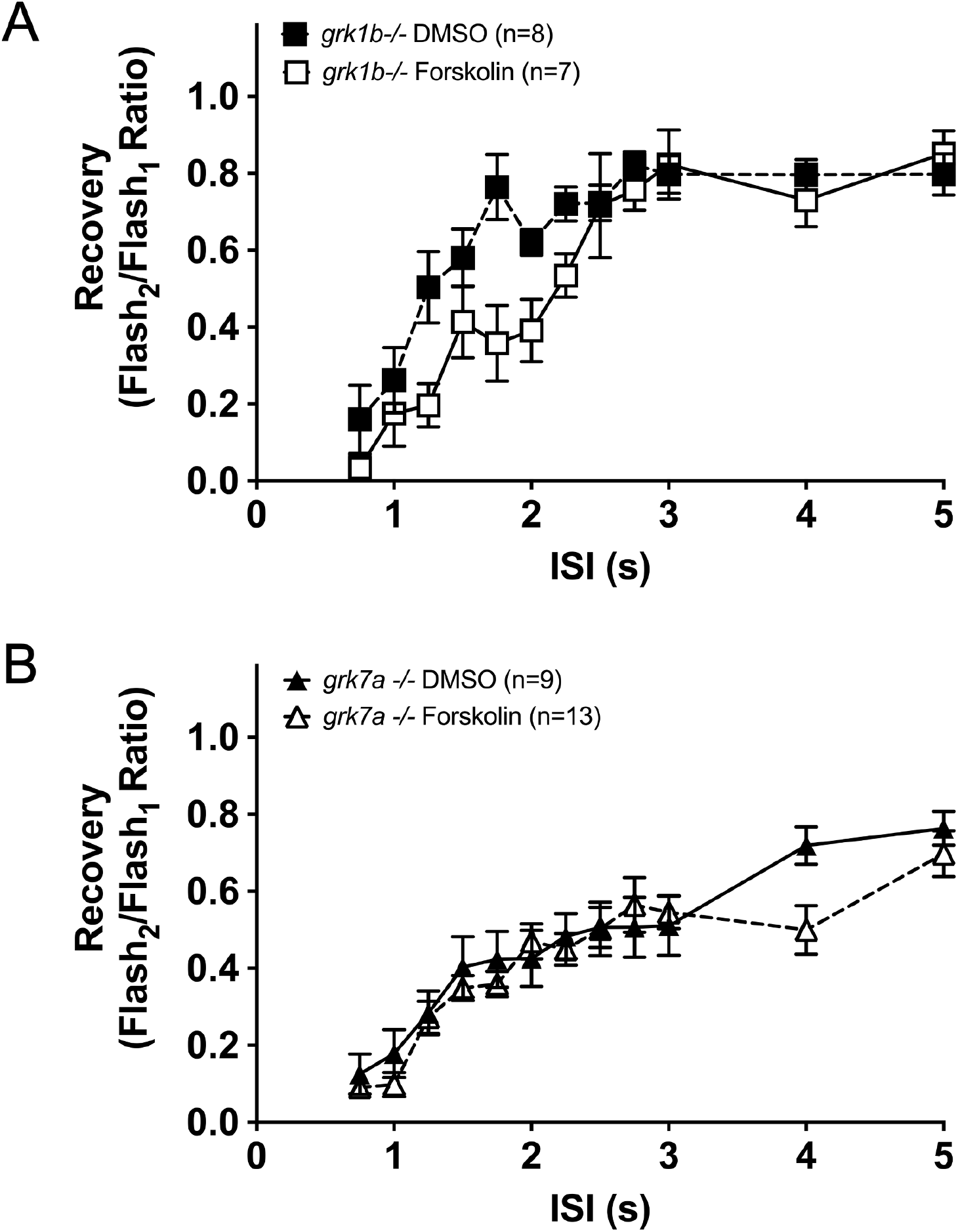
Recovery of the cone-mass receptor potential in Grk-knockout zebrafish larvae treated with forskolin. Treated larvae were subjected to successive stimuli using a dual flash paradigm of a 20-ms flash of saturating white light (1000 cd/m^2^) with an interstimulus interval (ISI) ranging from 0.75 s to 5 s. Cone-mass receptor potential recovery was plotted as the ratio of the maximum isolated cone mass receptor potential response of the second stimulus to that of the initial stimulus for A) *grk1b-/-* and B) *grk7a-/-* larvae. A linear mixed model analysis of covariance found a significant effect of forskolin when compared to vehicle for *grk1b-/-* [F_(1, 224)_ = 17.82; p <0.0001] larvae, but not *grk7a-/-* larvae [F_(1, 340)_ = 2.363; p=0.1252] Error bars represent SEM.

### Phosphorylation of Grk7 is mediated by PKA in vivo in zebrafish cones

While both Grk1 and Grk7 readily undergo phosphorylation by PKAα *in vitro*, the involvement of PKA in the phosphorylation of retinal GRKs *in vivo* is still speculative. Although PKA is a major downstream effector of cAMP, this cyclic nucleotide can activate signaling pathways through other effectors, such as Epac and cyclic nucleotide-gated ion channels (43,44). To determine the involvement of endogenous PKA in the phosphorylation of Grk7 in cones, transgenic *Tg(gnat2:prkar1a*^*G323D*^) zebrafish expressing a dominant negative RIα regulatory subunit (RIαB) that inhibits PKA activation and is driven by the cone transducin promotor (TαCP) were generated via tol2 transgenesis (30). After light-adaption for 2 hours, these larvae were dark adapted for 15 minutes and eyes were enucleated, homogenized, and subjected to immunoblot analysis. After 15 minutes of dark adaptation, levels of phosphorylated Grk7 were significantly decreased in fish expressing the PKA dominant negative in cone photoreceptors compared to wildtype fish (Figure 5B). This suggests that PKA is a critical effector in the phosphorylation of Grk7 in response to increased levels of cAMP in cones.

**Figure 5.**
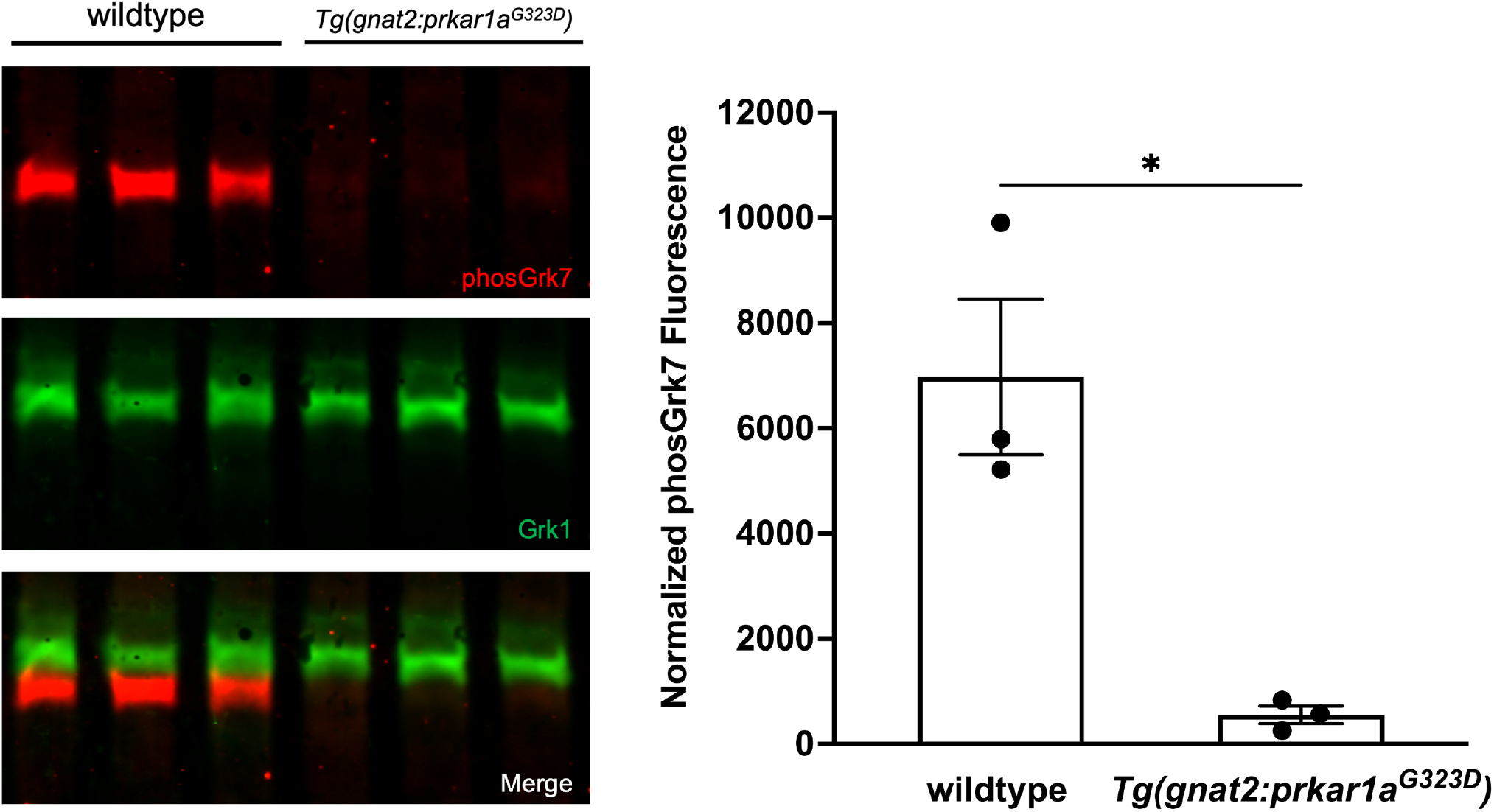
Dark-adapted Grk phosphorylation in 5 dpf transgenic *Tg(gnat2:prkar1a*^*G323D*^) zebrafish larvae expressing PKA dominant negative RIαB in cones. Larvae were dark-adapted for 15 minutes then euthanized and intact eyes were harvested for immunoblot analysis. Immunoblots were probed with antibodies against total Grk1 (green) and phosphorylated Grk7 (red) followed by incubation with secondary antibodies. The levels of detectable phosphorylated Grk7 were normalized against total Grk1 and quantified. Statistical comparison of multiple groups was performed using a Student’s *t* test. Error bars represent SEM (n=3). *P ≤ 0*.*05 (*)*.

### cAMP-dependent phosphorylation of Grk1 is undetectable in vertebrate cone photoreceptors

Our observation that Grk7a rather than Grk1 plays a greater role in modulating cone opsin lifetime *in vivo* in response to cAMP is consistent with observations from our previous report which showed that mice, which express only Grk1 in both rods and cones, display delayed dark adaptation in rods but not cones when Ser21 is mutated to block cAMP-dependent phosphorylation (45). In order determine whether cone-specific Grk1 is phosphorylated in response to changes in cAMP *in vivo*, the immunoblot analysis was modified. Using the previously characterized *grk1b-/-* fish along with a longer SDS-PAGE running period or a lower percentage acrylamide gel allows for specific identification of immunoblot bands corresponding to each zebrafish Grk1 paralog (Figure 6). Rod-expressed Grk1a (black arrowheads) migrates more slowly than cone-expressed Grk1b (black arrows), consistent with their predicted molecular weights of 64,185 Da and 63,567 Da, respectively. The lower band corresponding to Grk1b is undetectable in *grk1b-/-* fish and the ratio of Grk1a to Grk1b expression is higher in adult zebrafish (Figure 6B) than in larval zebrafish, most likely because the rod to cone ratio is higher in zebrafish adult retinas (46,47).

**Figure 6.**
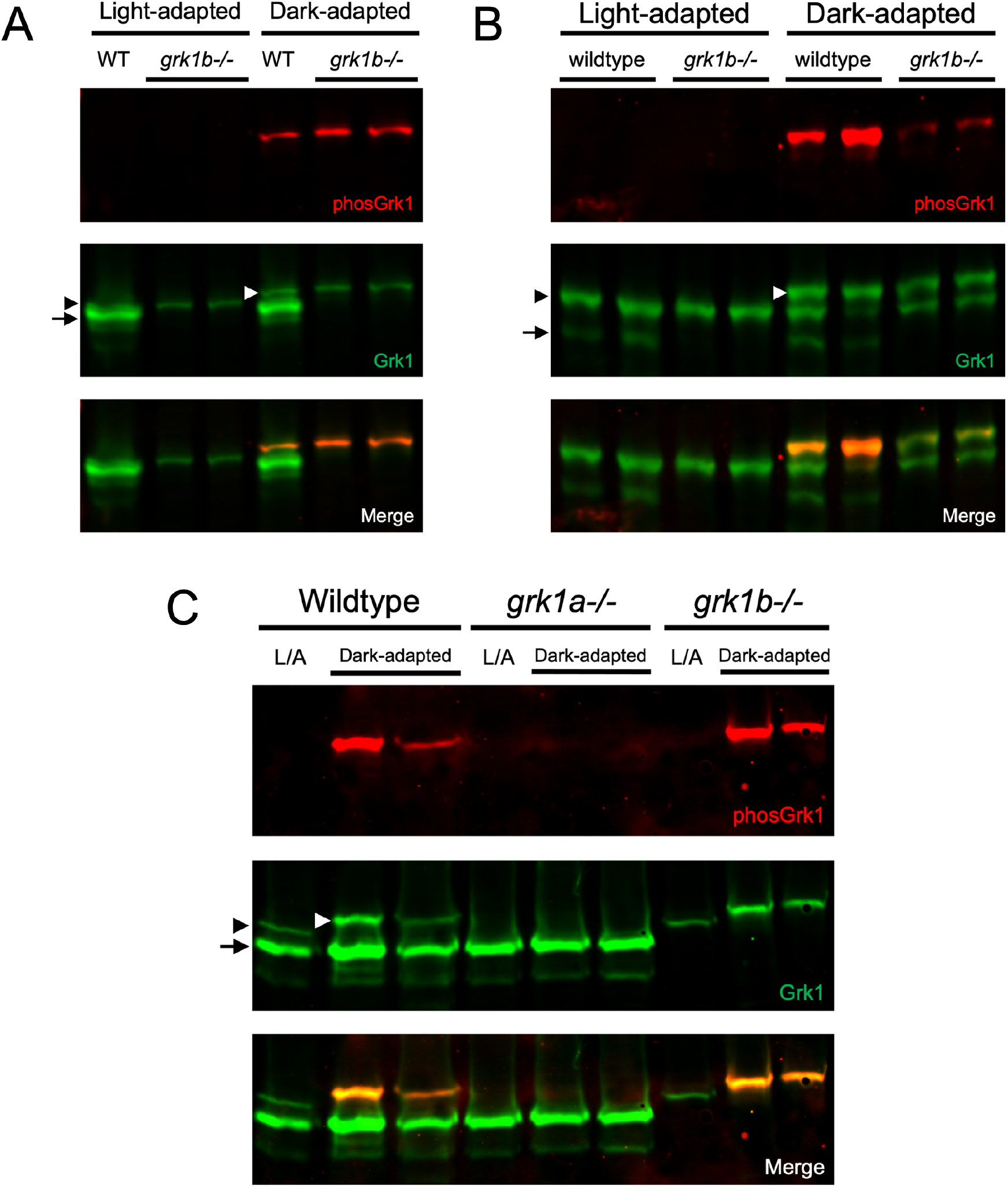
Grk1 phosphorylation in rod or cone Grk1 knockout zebrafish. Immunoblots were probed with antibodies against total Grk1 (green) and phosphorylated Grk1 (red) followed by incubation with secondary antibodies. Comparison of Grk1 phosphorylation in A) light- or dark-adapted wildtype and *grk1b-/-* larvae at 5 dpf, B) light- or dark-adapted wildtype and *grk1b-/-* adults, and C) light- or dark-adapted wildtype, *grk1b-/-*, and *grk1a-/-* larvae at 5 dpf. *Black arrowhead: Grk1a band; Black arrow: Grk1b band; White arrowhead, phosphorylated Grk1a band*.

Since the antibody against phosphorylated GRK1 does not discriminate between the Grk1a and Grk1b paralogs, and phosphorylated Grk1 (white arrowheads) is detectable in dark-adapted *grk1b-/-* larvae (Figure 6A) and adults (Figure 6B), it was unclear if both Grk1 paralogs are phosphorylated in a cAMP-dependent manner or just rod-expressed Grk1a. To determine if the phosphorylated Grk1 band detected in fish retinal homogenates is Grk1a, a *grk1a-/-* zebrafish (17bpdel95aa) was generated using CRISPR as described in Methods. Immunoblot analysis demonstrates that phosphorylated Grk1 is undetectable in dark-adapted *grk1a-/-* fish compared to wildtype and *grk1b-/-* zebrafish larvae (Fig. 6C). When *grk1b-/-* and *grk1a-/-* larvae were treated with forskolin, it was observed that *grk1b-/-* larvae exhibit a significant increase in Grk1 phosphorylation under both light-adapted and dark-adapted conditions (Figure 7A) whereas no significant Grk1 phosphorylation is detected in forskolin-treated *grk1a-/-* larvae (Figure 7B). Together, these data suggest that only rod-expressed Grk1a undergoes cAMP-dependent phosphorylation in zebrafish.

**Figure 7.**
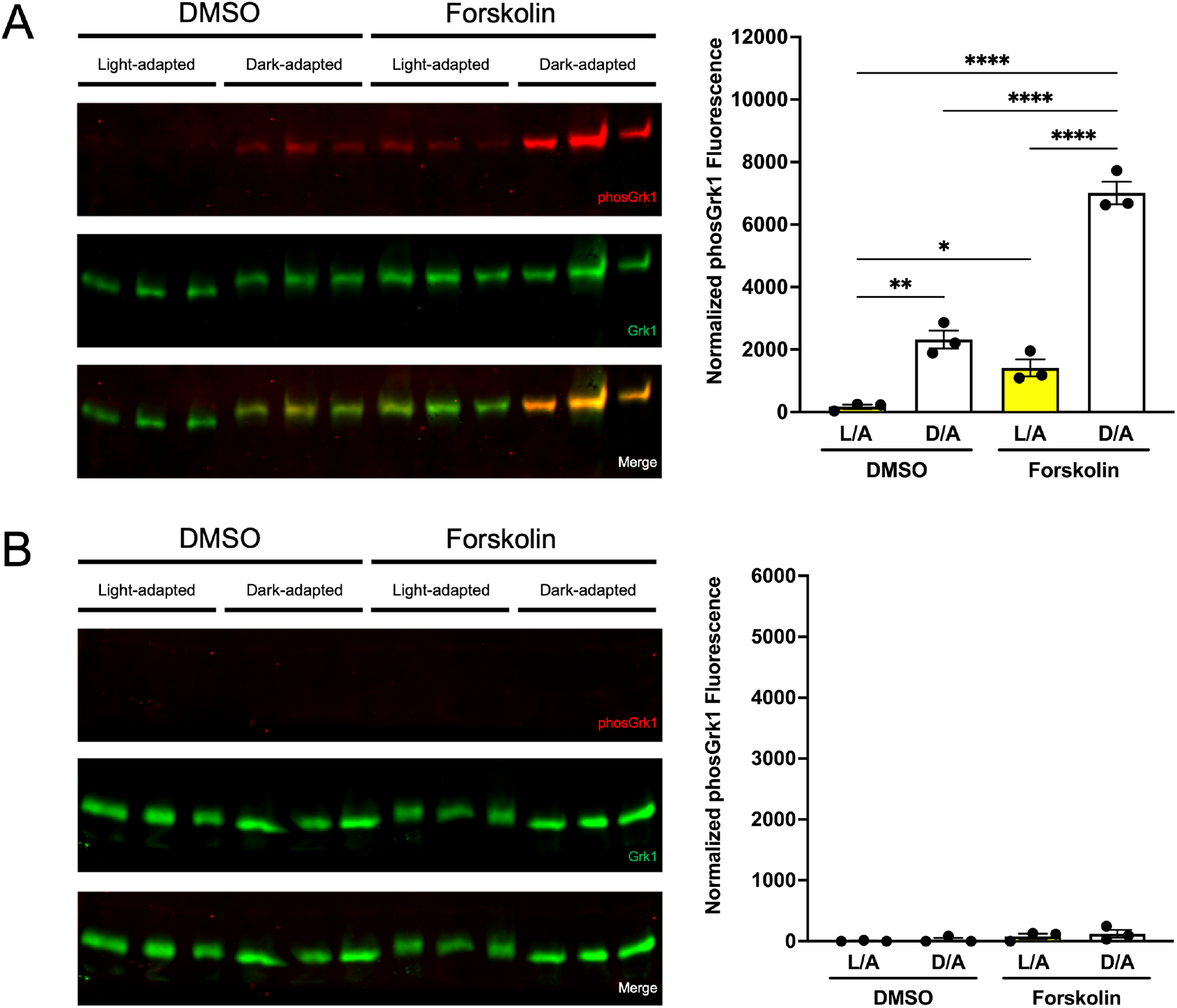
Effect of light exposure and forskolin incubation on phosphorylation of retinal GRKs in Grk1 knockout zebrafish larvae at 5 dpf. A) *grk1b-/-* or B) *grk1a-/-* larvae were light- or dark-adapted, followed by incubation with forskolin or DMSO for 30 minutes. Larvae were euthanized and intact eyes were harvested for immunoblot analysis. Immunoblots were probed with antibodies against total Grk1 (green) and phosphorylated Grk1 (red) followed by incubation with secondary antibodies. The levels of phosphorylated Grk1 were normalized against total Grk1 and quantified. Statistical comparison of multiple groups was performed using a two-way analysis of variance (ANOVA) followed by a Tukey post hoc test. Error bars represent SEM (n=3). *P ≤ 0*.*05 (*), P ≤ 0*.*01(**), P ≤ 0*.*001(***), P ≤ 0*.*0001 (****)*.

These observations are consistent with our previous report showing that mice with a mutation in Grk1 that converts the phosphorylation site, Ser21, to alanine have delayed dark adaptation in rods but not cones and led us to examine the phosphorylation of Grk1 in the ‘all-cone’ retina of *Nrl-/-* mice. Mice deficient for the Nrl transcription factor possess retinas lacking rods and instead consist mostly of modified S-cones (48). When immunoblots of retinal homogenates of light- or dark-adapted wildtype and *Nrl-/-* mice were compared, Grk1 was not phosphorylated in dark-adapted *Nrl-/-* mouse retinas compared to wildtype retinas (Figure 8). This suggests that a lack of cAMP-dependent Grk1 phosphorylation in cone photoreceptors may be intrinsic to vertebrates expressing Grk1 in cones.

**Figure 8.**
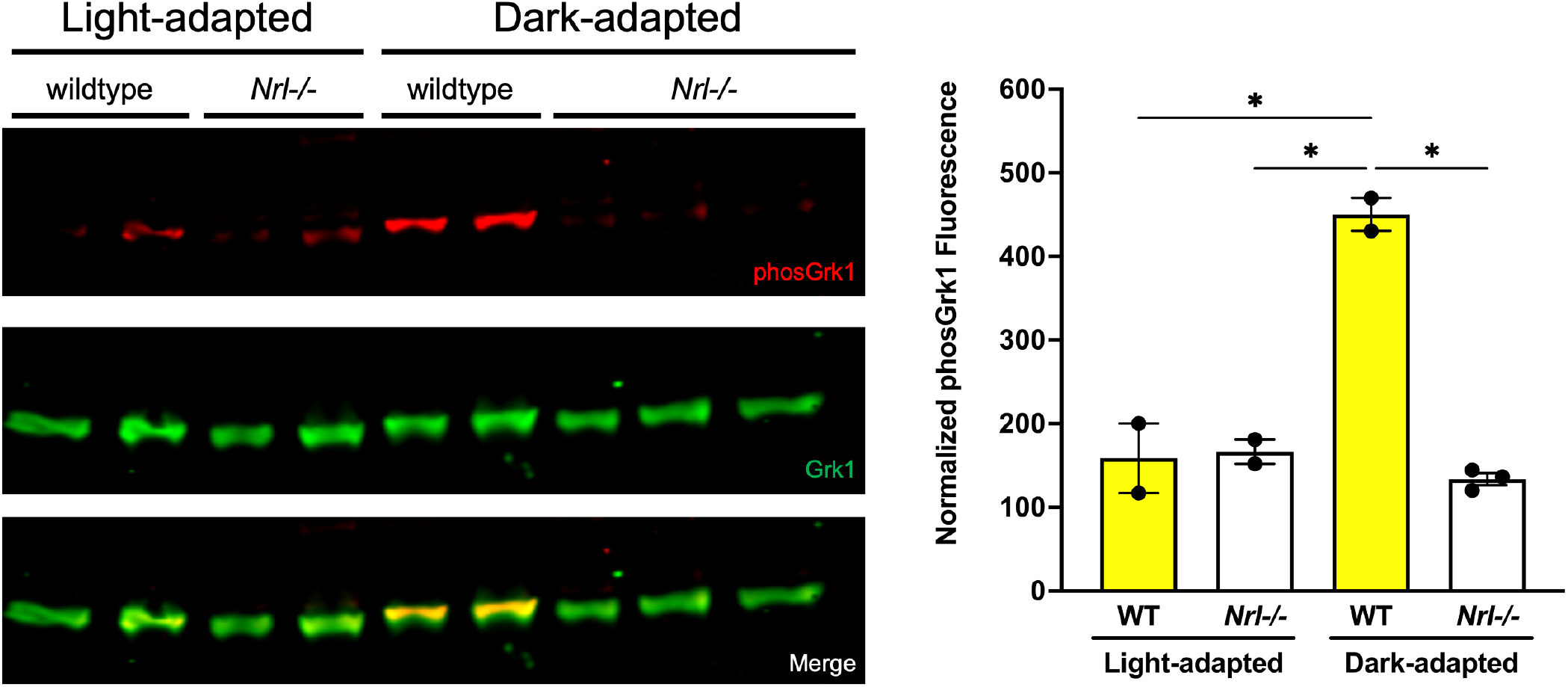
Grk1 phosphorylation in *Nrl-/-* mice. Adult mice were light- or dark-adapted, euthanized, and retinas harvested under the same light conditions. Immunoblots were probed with antibodies against total Grk1 (*green*) and phosphorylated Grk1 (*red*) followed by incubation with secondary antibodies. The levels of phosphorylated Grk1 were normalized against total Grk1 and quantified. Statistical comparison of multiple groups was performed using a two-way analysis of variance (ANOVA) followed by a Tukey post hoc test. Error bars represent SEM, (n=2-3). *P ≤ 0*.*05 (*)*

## Discussion

The present report demonstrates the *in vivo* phosphorylation of the photoreceptor GRKs in vertebrates, as well as what roles they may play in modulating cone phototransduction in the dark when intracellular cAMP levels are high. We chose zebrafish as our principal model, not only because they are an established model for the study of cone photoreceptor kinetics, but also because they have a similar cellular distribution profile of photoreceptor Grks as humans. In human retinas, Grk1 is expressed in both rods and cones while Grk7 expression is limited to cones (22). Zebrafish, as a result of a genome duplication event in teleost evolution, express one Grk1 ortholog (Grk1a) in rods and another (Grk1b) in cones, and a GRK7 ortholog (Grk7a) in cones (49–51). The phosphorylation of photoreceptor Grks *in vivo* is likely associated with the elevated levels of intracellular cAMP levels found in dark-adapted photoreceptors, of which PKA is a major downstream effector (18,19). Additionally, we previously reported that Grk1 phosphorylation is absent in dark-adapted mice when adenylyl cyclase expression is deleted and that *ex vivo* treatment of frog retinas with forskolin is associated with an increase in Grk7 phosphorylation independent of light conditions (16,17). Using an antibody against phosphorylated Grk7 and an antibody specific for phosphorylated zebrafish Grk1a and Grk1b, we demonstrate that levels of phosphorylated Grk1 and Grk7 are elevated in retinas of dark-adapted zebrafish similar to other species studied.

A study of frog rod photoreceptors treated with forskolin observed elevated levels of cAMP in the outer segments, while suction pipette recordings showed increased sensitivity of the rod photoresponse in a manner suggesting that multiple targets which could impact phototransduction are affected (41,52). This included an increase in PDE activity and a reduction in guanylate cyclase activity, as well as an increase in the Ca^2+^ exchanger current leading to an increase in intracellular Ca^2+^. Prior studies of increased intracellular Ca^2+^ in rods illustrated its ability to impact light adaptation through several targets such as increased affinity of recoverin for Grk1, decreased guanylyl cyclase activity via GCAPs, or a decreased channel affinity for cGMP via inhibition by Ca^2+^-calmodulin (53–59). Interestingly, a similar study of isolated carp cones treated with forskolin showed that while the rising and descending phases of the light response were slowed, flash sensitivity of carp cones were not affected in the same manner as rods (60). Using whole ERG analysis of intact larval zebrafish, we show that forskolin does not affect the normalized sensitivity of the cone photoresponse. However, forskolin is associated with a significant delay in recovery of the cone photoresponse to successive stimuli. Although increased intracellular cAMP likely has several downstream targets that impact the photoresponse, both our *in vitro* and *in vivo* observations of Grk phosphorylation led us to examine the effect of forskolin in zebrafish cone Grk knockout lines lacking either Grk1b or Grk7a. We found that while forskolin led to a decrease in cone photoresponse recovery in *grk1b-/-* larvae, there was no forskolin-associated decrease in cone recovery in *grk7a-/-*. This suggests that modulation of the cone photoresponse by elevated intracellular cAMP is mediated, at least in part, by Grk7a but not Grk1b.

Based on our previous observations that PKAα can phosphorylate GRKs *in vitro* and GRK phosphorylation *in vivo* coincides with elevated intracellular cAMP levels, it is reasonable to suggest that the retinal GRKs are phosphorylated by cAMP-activated PKA *in vivo*. The complete suppression of dark-adapted Grk7a phosphorylation in a zebrafish line expressing a dominant negative inhibitor of PKA in cones is the first proof that PKA plays such a role *in vivo*. While this shows a role for PKA in the phosphorylation of GRK7 *in vivo*, it does not rule out that GRK7 could be phosphorylated by a serine/threonine kinase activated downstream of PKA as has been shown for other proteins (61–63). Interestingly, while PKAα phosphorylated both GRK1 and GRK7 *in vitro*, our observations of Grk1 phosphorylation in retinas of *grk1a-/-* and *grk1b-/-* zebrafish as well as in the ‘all cone’ retinas of the *Nrl-/-* mouse strongly suggest that Grk1 in vertebrate cones is not phosphorylated in a cAMP-dependent manner. This observation agrees with both our previous report of GRK1-S21A mice displaying a significant delay in dark adaptation in rods but not cones, as well the results of this report showing that Grk7a but not Grk1b modulates recovery of the cone photoresponse in forskolin-treated zebrafish larvae.

The involvement of one or more downstream kinases with different specificities towards each GRK paralog or cellular expression profile could explain the differential phosphorylation of Grk1 between rod and cones photoreceptors, as well as that of Grk1 and Grk7 in cones. Alternatively, if PKA directly phosphorylates GRKs in photoreceptors, the differential phosphorylation could be the result of PKA having different affinities for each GRK paralog. A study using a fluorescent biosensor for PKA in mouse retinas showed that PKA was activated less efficiently in cones than in rods following the extinguishing of a short light stimulus (64), which might explain the differences in Grk1 phosphorylation between rods and cones. Another explanation for the lack of detectable cAMP-dependent Grk1 phosphorylation in cones could be a phosphatase that is likely cone-specific (to account for our observations in *Nrl-/-* mice) and which acts preferentially toward Grk1 rather than Grk7 (to account for our observations in zebrafish). Currently, the identity of any phosphatase that targets retinal GRKs is unknown.

Grk1 phosphorylation could also be sterically hindered in cones but not rods. Aside from its activity towards light-activated opsin, the most widely studied interaction with Grk1 has been that with recoverin. When intracellular Ca^2+^ levels are high, extrusion of a myristoyl switch from recoverin exposes a hydrophobic pocket that binds to the first 25 amino acids of Grk1 and blocks the interaction of the kinase with light-activated opsin (12–14). Considering that the same isoforms of recoverin and Grk1 are expressed in mammalian rods and cones, any potential steric inhibition of Grk1 phosphorylation by recoverin is likely to be associated with cone-specific differences in either recoverin expression levels or Ca^2+^ levels in response to light. In zebrafish photoreceptors, where multiple paralogs of recoverin are differentially expressed, there is greater likelihood that paralog-specific differences in recoverin-Grk1 interaction could account for Grk1 phosphorylation in rods but not in cones (65,66). Recent analysis using surface plasmon resonance has shown that zebrafish rcv1a, an ortholog of mouse recoverin that is expressed in rods and UV cones, appears to bind equally well to Grk1b in the presence or absence of Ca^2+^ (67). Studies of carp and frog photoreceptors showed that visinin, another neuronal calcium sensor in the same family as recoverin, was expressed at a much higher level in cones compared to recoverin in rods and had a slightly greater inhibition of Grk7 compared to Grk1 (68). Despite this, the same study reported that the level of inhibition of retinal Grks by visinin was indistinguishable from inhibition by recoverin (68). Additionally, the lack of cAMP-dependent phosphorylation of GRK1 in cones is also observed in mice which do not express visinin. It is also possible that an unidentified Grk1-interacting protein expressed in vertebrate cones could prevent phosphorylation in response to elevated intracellular cAMP.

In summary, our results show that intracellular cAMP levels modulate recovery of the cone photoresponse via PKA-mediated phosphorylation of Grk7 rather than Grk1 in zebrafish, a species with a retinal GRK expression profile similar to humans, and that cAMP-dependent phosphorylation of Grk1 is absent in cones in both zebrafish and mouse. These observations not only corroborate previous reports detailing the role of cAMP-dependent phosphorylation of retina GRKS *in vivo* and *in vitro*, but will help us to better understand the mechanisms of cone adaptation and recovery in relation to human photoreceptor biology.

## Acknowledgments

The authors thank Jessica Gumerson & Anand Swaroop of the Neurobiology Neurodegeneration & Repair Laboratory (NNRL) at the National Eye Institute for the collection and shipment of light-adapted and dark-adapted *Nrl-/-* mouse retinas. We would also like to thank Michelle Altemara, MS, and the Zebrafish Aquaculture Core Facility at The University of North Carolina at Chapel Hill for maintenance of zebrafish stocks.

Supported by National Institutes of Health Grants NEI R01EY012224 (ERW).

Disclosure: J.D. Chrispell, None; Y. Xiong, None; E.R. Weiss, None.

## Works Cited

1. Hecht, S., Shlaer, S., and Pirenne, M. H. (1942) Energy, Quanta, and Vision. J Gen Physiol 25, 819–840

2. Baylor, D. A. (1987) Photoreceptor signals and vision. Proctor lecture. Invest Ophthalmol Vis Sci 28, 34–49

3. Yau, K. W. (1994) Phototransduction mechanism in retinal rods and cones. The Friedenwald Lecture. Invest Ophthalmol Vis Sci 35, 9–32

4. Arshavsky, V. Y., and Burns, M. E. (2012) Photoreceptor signaling: supporting vision across a wide range of light intensities. J Biol Chem 287, 1620–1626

5. Latek, D., Modzelewska, A., Trzaskowski, B., Palczewski, K., and Filipek, S. (2012) G protein-coupled receptors--recent advances. Acta Biochim Pol 59, 515–529

6. Gross, O. P., and Burns, M. E. (2010) Control of rhodopsin’s active lifetime by arrestin-1 expression in mammalian rods. J Neurosci 30, 3450–3457

7. Zhao, X., Huang, J., Khani, S. C., and Palczewski, K. (1998) Molecular forms of human rhodopsin kinase (GRK1). J Biol Chem 273, 5124–5131

8. Kuhn, H., and Wilden, U. (1987) Deactivation of photoactivated rhodopsin by rhodopsin-kinase and arrestin. J Recept Res 7, 283–298

9. Chen, C. K., Zhang, K., Church-Kopish, J., Huang, W., Zhang, H., Chen, Y. J., Frederick, J. M., and Baehr, W. (2001) Characterization of human GRK7 as a potential cone opsin kinase. Mol Vis 7, 305–313

10. Maeda, T., Imanishi, Y., and Palczewski, K. (2003) Rhodopsin phosphorylation: 30 years later. Prog Retin Eye Res 22, 417–434

11. Tachibanaki, S., Arinobu, D., Shimauchi-Matsukawa, Y., Tsushima, S., and Kawamura, S. (2005) Highly effective phosphorylation by G protein-coupled receptor kinase 7 of light-activated visual pigment in cones. Proc Natl Acad Sci U S A 102, 9329–9334

12. Ames, J. B., Tanaka, T., Ikura, M., and Stryer, L. (1995) Nuclear magnetic resonance evidence for Ca(2+)-induced extrusion of the myristoyl group of recoverin. J Biol Chem 270, 30909–30913

13. Ames, J. B., Ishima, R., Tanaka, T., Gordon, J. I., Stryer, L., and Ikura, M. (1997) Molecular mechanics of calcium-myristoyl switches. Nature 389, 198–202

14. Ames, J. B., Levay, K., Wingard, J. N., Lusin, J. D., and Slepak, V. Z. (2006) Structural basis for calcium-induced inhibition of rhodopsin kinase by recoverin. J Biol Chem 281, 37237–37245

15. Horner, T. J., Osawa, S., Schaller, M. D., and Weiss, E. R. (2005) Phosphorylation of GRK1 and GRK7 by cAMP-dependent protein kinase attenuates their enzymatic activities. J Biol Chem 280, 28241–28250

16. Osawa, S., Jo, R., and Weiss, E. R. (2008) Phosphorylation of GRK7 by PKA in cone photoreceptor cells is regulated by light. J Neurochem 107, 1314–1324

17. Osawa, S., Jo, R., Xiong, Y., Reidel, B., Tserentsoodol, N., Arshavsky, V. Y., Iuvone, P. M., and Weiss, E. R. (2011) Phosphorylation of G protein-coupled receptor kinase 1 (GRK1) is regulated by light but independent of phototransduction in rod photoreceptors. J Biol Chem 286, 20923–20929

18. Farber, D. B., Souza, D. W., Chase, D. G., and Lolley, R. N. (1981) Cyclic nucleotides of cone-dominant retinas. Reduction of cyclic AMP levels by light and by cone degeneration. Invest Ophthalmol Vis Sci 20, 24–31

19. Cohen, A. I., and Blazynski, C. (1990) Dopamine and its agonists reduce a light-sensitive pool of cyclic AMP in mouse photoreceptors. Vis Neurosci 4, 43–52

20. Weiss, E. R., Hao, Y., Dickerson, C. D., Osawa, S., Shi, W., Zhang, L., and Wong, F. (1995) Altered cAMP levels in retinas from transgenic mice expressing a rhodopsin mutant. Biochem Biophys Res Commun 216, 755–761

21. Traverso, V., Bush, R. A., Sieving, P. A., and Deretic, D. (2002) Retinal cAMP levels during the progression of retinal degeneration in rhodopsin P23H and S334ter transgenic rats. Invest Ophthalmol Vis Sci 43, 1655–1661

22. Weiss, E. R., Ducceschi, M. H., Horner, T. J., Li, A., Craft, C. M., and Osawa, S. (2001) Species-specific differences in expression of G-protein-coupled receptor kinase (GRK) 7 and GRK1 in mammalian cone photoreceptor cells: implications for cone cell phototransduction. J Neurosci 21, 9175–9184

23. Wada, Y., Sugiyama, J., Okano, T., and Fukada, Y. (2006) GRK1 and GRK7: unique cellular distribution and widely different activities of opsin phosphorylation in the zebrafish rods and cones. J Neurochem 98, 824–837

24. Fuchs, S., Nakazawa, M., Maw, M., Tamai, M., Oguchi, Y., and Gal, A. (1995) A homozygous 1-base pair deletion in the arrestin gene is a frequent cause of Oguchi disease in Japanese. Nat Genet 10, 360–362

25. Yamamoto, S., Sippel, K. C., Berson, E. L., and Dryja, T. P. (1997) Defects in the rhodopsin kinase gene in the Oguchi form of stationary night blindness. Nat Genet 15, 175–178

26. Cideciyan, A. V., Jacobson, S. G., Gupta, N., Osawa, S., Locke, K. G., Weiss, E. R., Wright, A. F., Birch, D. G., and Milam, A. H. (2003) Cone deactivation kinetics and GRK1/GRK7 expression in enhanced S cone syndrome caused by mutations in NR2E3. Invest Ophthalmol Vis Sci 44, 1268–1274

27. Chrispell, J. D., Dong, E., Osawa, S., Liu, J., Cameron, D. J., and Weiss, E. R. (2018) Grk1b and Grk7a Both Contribute to the Recovery of the Isolated Cone Photoresponse in Larval Zebrafish. Invest Ophthalmol Vis Sci 59, 5116–5124

28. Kolesnikov, A. V., Chrispell, J. D., Osawa, S., Kefalov, V. J., and Weiss, E. R. (2020) Phosphorylation at Serine 21 in G protein-coupled receptor kinase 1 (GRK1) is required for normal kinetics of dark adaption in rod but not cone photoreceptors. FASEB J 34, 2677–2690

29. Hwang, W. Y., Fu, Y., Reyon, D., Maeder, M. L., Tsai, S. Q., Sander, J. D., Peterson, R. T., Yeh, J. R., and Joung, J. K. (2013) Efficient genome editing in zebrafish using a CRISPR-Cas system. Nat Biotechnol 31, 227–229

30. Willis, B. S., Niswender, C. M., Su, T., Amieux, P. S., and McKnight, G. S. (2011) Cell-type specific expression of a dominant negative PKA mutation in mice. PLoS One 6, e18772

31. Woodford, T. A., Correll, L. A., McKnight, G. S., and Corbin, J. D. (1989) Expression and characterization of mutant forms of the type I regulatory subunit of cAMP-dependent protein kinase. The effect of defective cAMP binding on holoenzyme activation. J Biol Chem 264, 13321–13328

32. Liu, H., and Naismith, J. H. (2008) An efficient one-step site-directed deletion, insertion, single and multiple-site plasmid mutagenesis protocol. BMC Biotechnol 8, 91

33. Kwan, K. M., Fujimoto, E., Grabher, C., Mangum, B. D., Hardy, M. E., Campbell, D. S., Parant, J. M., Yost, H. J., Kanki, J. P., and Chien, C. B. (2007) The Tol2kit: a multisite gateway-based construction kit for Tol2 transposon transgenesis constructs. Dev Dyn 236, 3088–3099

34. Kennedy, B. N., Alvarez, Y., Brockerhoff, S. E., Stearns, G. W., Sapetto-Rebow, B., Taylor, M. R., and Hurley, J. B. (2007) Identification of a zebrafish cone photoreceptor-specific promoter and genetic rescue of achromatopsia in the nof mutant. Invest Ophthalmol Vis Sci 48, 522–529

35. Lee, B. Y., Thulin, C. D., and Willardson, B. M. (2004) Site-specific phosphorylation of phosducin in intact retina. Dynamics of phosphorylation and effects on G protein beta gamma dimer binding. J Biol Chem 279, 54008–54017

36. Laemmli, U. K. (1970) Cleavage of structural proteins during the assembly of the head of bacteriophage T4. Nature 227, 680–685

37. Korenbrot, J. I., Mehta, M., Tserentsoodol, N., Postlethwait, J. H., and Rebrik, T. I. (2013) EML1 (CNG-modulin) controls light sensitivity in darkness and under continuous illumination in zebrafish retinal cone photoreceptors. J Neurosci 33, 17763–17776

38. Chrispell, J. D., Rebrik, T. I., and Weiss, E. R. (2015) Electroretinogram analysis of the visual response in zebrafish larvae. J Vis Exp

39. Kim, D. Y., and Jung, C. S. (2012) Gap junction contributions to the goldfish electroretinogram at the photopic illumination level. Korean J Physiol Pharmacol 16, 219–224

40. Naka, K. I., and Rushton, W. A. (1966) S-potentials from colour units in the retina of fish (Cyprinidae). J Physiol 185, 536–555

41. Astakhova, L. A., Samoiliuk, E. V., Govardovskii, V. I., and Firsov, M. L. (2012) cAMP controls rod photoreceptor sensitivity via multiple targets in the phototransduction cascade. J Gen Physiol 140, 421–433

42. Nowak, J. Z., Sek, B., and Zurawska, E. (1990) Activation of D2 dopamine receptors in hen retina decreases forskolin-stimulated cyclic AMP accumulation and serotonin N-acetyltransferase (NAT) activity. Neurochem Int 16, 73–80

43. Manoury, B., Idres, S., Leblais, V., and Fischmeister, R. (2020) Ion channels as effectors of cyclic nucleotide pathways: Functional relevance for arterial tone regulation. Pharmacol Ther 209, 107499

44. Bos, J. L. (2003) Epac: a new cAMP target and new avenues in cAMP research. Nat Rev Mol Cell Biol 4, 733–738

45. Kolesnikov, A. V., Chrispell, J., Kefalov, V., and Weiss, E. R. (2019) Ablation of cAMP-dependent GRK1 phosphorylation suppresses dark adaption of rod photoreceptors. Invest Ophth Vis Sci 60

46. Johns, P. R. (1982) Formation of photoreceptors in larval and adult goldfish. J Neurosci 2, 178–198

47. Fadool, J. M. (2003) Development of a rod photoreceptor mosaic revealed in transgenic zebrafish. Dev Biol 258, 277–290

48. Mears, A. J., Kondo, M., Swain, P. K., Takada, Y., Bush, R. A., Saunders, T. L., Sieving, P. A., and Swaroop, A. (2001) Nrl is required for rod photoreceptor development. Nat Genet 29, 447–452

49. Volff, J. N. (2005) Genome evolution and biodiversity in teleost fish. Heredity (Edinb) 94, 280–294

50. Rinner, O., Makhankov, Y. V., Biehlmaier, O., and Neuhauss, S. C. (2005) Knockdown of cone-specific kinase GRK7 in larval zebrafish leads to impaired cone response recovery and delayed dark adaptation. Neuron 47, 231–242

51. Makhankov, Y. V., Rinner, O., and Neuhauss, S. C. (2004) An inexpensive device for non-invasive electroretinography in small aquatic vertebrates. J Neurosci Methods 135, 205–210

52. Astakhova, L. A., Nikolaeva, D. A., Fedotkina, T. V., Govardovskii, V. I., and Firsov, M. L. (2017) Elevated cAMP improves signal-to-noise ratio in amphibian rod photoreceptors. J Gen Physiol 149, 689–701

53. Chen, C. K., Inglese, J., Lefkowitz, R. J., and Hurley, J. B. (1995) Ca(2+)-dependent interaction of recoverin with rhodopsin kinase. J Biol Chem 270, 18060–18066

54. Palczewski, K., Polans, A. S., Baehr, W., and Ames, J. B. (2000) Ca(2+)-binding proteins in the retina: structure, function, and the etiology of human visual diseases. Bioessays 22, 337–350

55. Burns, M. E., Mendez, A., Chen, J., and Baylor, D. A. (2002) Dynamics of cyclic GMP synthesis in retinal rods. Neuron 36, 81–91

56. Dizhoor, A. M., Olshevskaya, E. V., and Peshenko, I. V. (2010) Mg2+/Ca2+ cation binding cycle of guanylyl cyclase activating proteins (GCAPs): role in regulation of photoreceptor guanylyl cyclase. Mol Cell Biochem 334, 117–124

57. Koutalos, Y., Nakatani, K., Tamura, T., and Yau, K. W. (1995) Characterization of guanylate cyclase activity in single retinal rod outer segments. J Gen Physiol 106, 863–890

58. Koutalos, Y., Nakatani, K., and Yau, K. W. (1995) The cGMP-phosphodiesterase and its contribution to sensitivity regulation in retinal rods. J Gen Physiol 106, 891–921

59. Chen, J., Woodruff, M. L., Wang, T., Concepcion, F. A., Tranchina, D., and Fain, G. L. (2010) Channel modulation and the mechanism of light adaptation in mouse rods. J Neurosci 30, 16232–16240

60. Sitnikova, V. S., Astakhova, L. A., and Firsov, M. L. (2021) cAMP-Dependent Regulation of the Phototransduction Cascade in Cones. Neuroscience and Behavioral Physiology 51, 108–115

61. Ko, G. Y., Ko, M., and Dryer, S. E. (2004) Circadian and cAMP-dependent modulation of retinal cone cGMP-gated channels does not require protein synthesis or calcium influx through L-type channels. Brain Res 1021, 277–280

62. Ko, G. Y., Ko, M. L., and Dryer, S. E. (2004) Circadian regulation of cGMP-gated channels of vertebrate cone photoreceptors: role of cAMP and Ras. J Neurosci 24, 1296–1304

63. Isobe, K., Jung, H. J., Yang, C. R., Claxton, J., Sandoval, P., Burg, M. B., Raghuram, V., and Knepper, M. A. (2017) Systems-level identification of PKA-dependent signaling in epithelial cells. Proc Natl Acad Sci U S A 114, E8875–E8884

64. Sato, S., Yamashita, T., and Matsuda, M. (2020) Rhodopsin-mediated light-off-induced protein kinase A activation in mouse rod photoreceptor cells. Proc Natl Acad Sci U S A 117, 26996–27003

65. Zang, J., Keim, J., Kastenhuber, E., Gesemann, M., and Neuhauss, S. C. (2015) Recoverin depletion accelerates cone photoresponse recovery. Open Biol 5

66. Zang, J., and Neuhauss, S. C. F. (2018) The Binding Properties and Physiological Functions of Recoverin. Front Mol Neurosci 11, 473

67. Ahrens, N., Elbers, D., Greb, H., Janssen-Bienhold, U., and Koch, K. W. (2021) Interaction of G protein-coupled receptor kinases and recoverin isoforms is determined by localization in zebrafish photoreceptors. Biochim Biophys Acta Mol Cell Res 1868, 118946

68. Arinobu, D., Tachibanaki, S., and Kawamura, S. (2010) Larger inhibition of visual pigment kinase in cones than in rods. J Neurochem 115, 259–268

